# Contribution of FEF to attentional periodicity during visual search: a TMS study

**DOI:** 10.1101/414383

**Authors:** Laura Dugué, Alexy Asaf Beck, Philippe Marque, Rufin VanRullen

## Abstract

Visual search, looking for a target embedded among distractors, has long been used to study attention. Current theories postulate a two-stage process in which early visual areas perform feature extraction, while higher-order regions perform attentional selection. Such a model implies iterative communication between low- and high-level regions to sequentially select candidate targets in the array, focus attention on these elements, and eventually permit target recognition. This leads to two predictions: (1) high-level, attentional regions and (2) early visual regions should both be iteratively (periodically) involved during the search. Here, we used Transcranial Magnetic Stimulation (TMS) applied over the Frontal-Eye Field (FEF), known to be involved in attentional selection, at various delays while observers performed a difficult, attentional search task. We observed a periodic pattern of interference at 7 Hz (theta) suggesting that the FEF is periodically involved during this difficult search task. We further compared this result with two previous studies (Dugué et al., 2011; 2015a) in which a similar TMS procedure was applied over the early visual cortex (V1) while observers performed the same task. This analysis revealed, for both studies, the same pattern of interference, i.e. V1 is periodically involved during this difficult search task, at the theta frequency. Together, these converging findings confirm our predictions that difficult search is supported by the periodic involvement of both low- and high-level regions, at the theta frequency.

**Significant statement:** Attention models postulate a two-stage process during visual search in which early visual regions perform feature extraction, while higher-order regions perform attentional selection, these two levels iteratively (periodically) communicating until target recognition. Using TMS, we tested whether there is a causal link between both attentional and early visual regions, and attentional search performance. We showed that a difficult, attentional search is supported by the periodic involvement of both V1 and the FEF, at the theta frequency (∼6-7 Hz). This finding support the idea that visual search tasks are processed by a hierarchical system involving periodic, iterative connections between low- and high-level regions allowing successful attentional exploration.

## Introduction

Covert attention selectively enhances visual processing at the attended location in the absence of eye movement. In the past decade, researchers studying the temporal dynamics of visual information processing have proposed that attention samples visual information periodically at low frequencies, theta (5-7 Hz; VanRullen et al., 2007; Busch and VanRullen, 2010; Landau and Fries, 2012; Fiebelkorn et al., 2013; VanRullen, 2013; Song et al., 2014; Huang et al., 2015; Landau et al., 2015; Dugué et al., 2015a; 2015b; 2016; 2017; Fiebelkorn et al., 2018; Helfrich et al., 2018) and alpha (8-12 Hz; Dugué and VanRullen, 2014; van Diepen et al., 2016; see VanRullen, 2016, for a comprehensive review). Critically, it has been proposed that the distinction between theta and alpha periodicity comes from the spatial exploration of the visual scene by attention (Dugué and VanRullen, 2017). In other words, when attention is not critical for the task, visual information is processed at the alpha frequency, while when attention explores the visual space (e.g. in cueing or visual search tasks), then visual information is processed at the theta frequency.

Visual search tasks, in which observers look for a target embedded among distractors, have long been used to study attentional deployment (see reviews: Eckstein, 2011; Nakayama and Martini, 2011). In search tasks known as difficult, authors have proposed a hierarchical processing stream (Palmer et al., 1993; Treisman, 1998; Itti and Koch, 2001; Deco et al., 2002). An early stage, presumably supported by early visual areas, would decompose the visual scene in given features (e.g. color, orientation, etc.). A high-level stage would then perform attentional selection, i.e. a priority map would select the spatial location of a candidate target to focus attentional resources on. The facilitation of target processing would then occur by sending feedback connections (Juan and Walsh, 2003; Saalmann et al., 2007; Dugué et al., 2011; 2015a) to the corresponding retinotopic region (Motter, 1994; Mehta et al., 2000; Schroeder et al., 2001; Kastner and Pinsk, 2004; Bressler et al., 2008). In such a model, this selection would iterate until target recognition, suggesting that both high-level, attentional regions and early visual regions would both be iteratively (periodically) involved during the search (Dugué et al., 2015a; Dugué and VanRullen, 2017). This is the prediction we tested here.

We used Transcranial Magnetic Stimulation (TMS) applied over the Frontal-Eye Field (FEF), known to be involved in attentional selection (Kastner et al., 1999; Corbetta and Shulman, 2002), at various delays while observers performed a difficult, attentional search task. We compared the results to two previously published studies (Dugué et al., 2011; 2015a) using the same difficult search task (finding the letter T among L letters) while observers were stimulated over the occipital pole (V1/V2) using a similar TMS protocol (see meta-analysis in **Table 1**). We found that both the FEF and early visual cortex were periodically involved during the difficult search task, at the theta frequency (∼6-7 Hz).

**Table 1:** TMS studies investigating the role of attention during difficult, visual search tasks. For each study we report the behavioral manipulation, the stimulation parameters: tested region (R: Right; L: Left; FEF: Frontal Eye Field; V5: MT area; PC: Parietal Cortex; PPC: Posterior Parietal Cortex; AG: Angular Gyrus, part of the PPC; STG: Superior Temporal Gyrus; DLPFC: Dorsolateral prefrontal cortex, right hemisphere), control condition, type of stimulation (TBS: Theta Burst Stimulation; note: otherwise mentioned, the reference is the onset of the visual stimuli) and the intensity of the stimulation (MSI: Maximum output Stimulation Intensity of the TMS machine), the amount of participants for which data were analyzed (n), and finally, the authors’ conclusions.

| Study | Behavioral manipulation | Stimulation parameters |  |  |  | n | Conclusions |
| --- | --- | --- | --- | --- | --- | --- | --- |
|  |  | Test Region | Control | Type of stimulation | Stimulation intensity |  |  |
| (Ashbridge, 1997) | Feature: color<br>Conjunction: color-orientation | R-PC | no-TMS | Single-pulse at 11 possible delays (from 0 to 200 ms) | 80% MSI | 5 | R-PC involved in conjunction but not feature tasks |
| (Walsh et al., 1998) | Conjunction: color-orientation | R-PC | no-TMS | Single-pulse at 11 possible delays (from 0 to 200 ms) | 80% MSI | 3 | R-PC involved in novel but not learned conjunction tasks |
| (Juan and Walsh, 2003) | Feature: color<br>Conjunction: color-orientation | V1/V2 | no-TMS | 10Hz for 500ms & Double-pulse (40 or 100ms interval) at 6 possible delays | 65% MSI | 8, 6 & 6 | V1 involved at late delays (feedback) during conjunction but not feature tasks |
| (Muggleton et al., 2003) | Feature: color<br>Conjunction: color-orientation | R-FEF<br>L-FEF | Vertex<br>V5 | 10Hz for 500ms | 65% MSI | 5 | FEF is involved in visual selection, in the absence of saccade |
| (Ellison et al., 2004) | Feature: orientation<br>Conjunction: color-orientation | R-PPC<br>R-STG | SHAM on<br>R-PPC or<br>R-STG | R-PPC: 10Hz for 500ms<br>R-STG: 4Hz for 500ms | 65% MSI | 5 | R-PPC involved in conjunction and R-STG in feature |
| (O'Shea et al., 2004) | Conjunction:<br>color-orientation | R-FEF | Vertex<br>V5 | Double-pulse<br>(40ms interval) at 5<br>possible delays<br>(from 0 to 120 ms),<br>40ms before mask<br>onset | FEF &<br>Vertex:<br>65% MSI<br>V5: 110%<br>phosphene<br>threshold | 12 | FEF is involved in<br>visual discrimination,<br>in the absence of<br>saccade |
| (Fuggetta et al., 2006) | Conjunction:<br>color-orientation | R-PPC | Vertex | Single-pulse 100ms<br>after stimulus onset | 85% MSI<br>on average | 7 | R-PPC involved in<br>conjunction task |
| (Ellison et al., 2007) | Conjunction:<br>color-orientation<br>and motion-<br>orientation | R-PPC<br>R-V5 | SHAM on<br>R-PPC or<br>R-V5 | 10Hz for 500ms | 65% MSI | 7 | R-PPC involved in<br>color-orientation<br>R-V5 in motion-<br>orientation |
| (Kalla et al., 2008) | Conjunction:<br>color-orientation | R-FEF<br>R-PPC | no-TMS | Double-pulse<br>(40ms interval) at 5<br>possible delays<br>(from 0 to 200 ms) | 60% MSI | 9 | R-FEF involved<br>earlier than R-PPC in<br>conjunction tasks |
| (Muggleton et al., 2008) | Feature: color<br>Conjunction:<br>color-orientation | R-AG<br>L-AG | Vertex<br>no-TMS | 10Hz for 500ms | 65% MSI | 8 | R-AG but not L-AG<br>involved in<br>conjunction but not<br>feature tasks |
| (Schindler et al., 2008) | Feature:<br>orientation<br>Conjunction:<br>color-orientation | R-PPC<br>R-STG | SHAM on<br>R-PPC or<br>R-STG | R-PPC: 10Hz for<br>500ms<br>R-STG: 4Hz for<br>500ms | 65% MSI | 5 | R-STG processes<br>the left part of the<br>search array<br>presented<br>contralateral<br>R-PPC involved for<br>left visual field when<br>full-fields array |
| (Kalla et al., 2009) | Feature: color<br>Conjunction:<br>color-orientation | DLPFC | Vertex<br>V5 | TBS: 50Hz bursts<br>at 5Hz | 40% MSI | 12 | DLPCF involved in<br>conjunction but not<br>feature tasks |
| (Dugué et al., 2011) | Conjunction:<br>L vs. T and L vs.<br>+ | V1/V2 | Stimulus<br>at non-<br>retinotopic<br>location | Double-pulse<br>(25ms interval) at 8<br>possible delays<br>(from 100 to 450<br>ms) | 55% MSI | 11 | V1 involved at late<br>delays (feedback)<br>during conjunction<br>but not feature tasks |
| (Lane et al., 2011) | Feature:<br>shape<br>Conjunction:<br>color-orientation | R-PPC | SHAM on<br>R-PPC | 10Hz for 500ms | 65% MSI | 12 | R-PPC involved in<br>conjunction and<br>feature tasks when<br>participants have to<br>point to the target,<br>but only conjunction<br>when button press<br>response |
| (Muggleton et al., 2011) | Conjunction:<br>color-orientation | R-FEF<br>L-PPC | no-TMS | 10Hz for 500ms | 60% MSI | 8 | L-PPC involved<br>when manual motor<br>response required |
| (Dugué et al., 2015a) | Conjunction:<br>L vs. T and L vs.<br>+ | V1/V2 | Stimulus<br>at non-<br>retinotopic<br>location | Double-pulse: one<br>at 312.5 ms and<br>one 13 possible<br>delays (from 112.5<br>to 437.5 ms) | 55% MSI | 10 | V1 involved<br>periodically during<br>conjunction but not<br>feature tasks (5hz) |
| (Yan, 2016) | Feature:<br>color<br>Conjunction:<br>orientation of<br>triangles | R-DLPFC<br>R-PPC | Vertex on<br>another<br>group of<br>participant<br>s | Double-pulse<br>(100ms interval)<br>200ms before<br>search array onset | 51% MSI<br>on average | 16 | DLPFC is involved in<br>the conjunction<br>search, while the<br>PPC is involved in<br>the feature search |
| Present<br>study | Conjunction:<br>L vs. T | R-FEF | Vertex | Double-pulse<br>(25ms interval) at 9<br>possible delays<br>(from 50 to 450 ms) | 50% MSI | 21 | FEF involved<br>periodically during a<br>conjunction task<br>(7hz) |

### Material and Methods

#### Participants

Twenty-three participants (7 women), aged 24-38 years old, were recruited. Two did not complete the experiment because of discomfort due to the stimulation. All participants gave written informed consent before the experiment. Standard exclusion criteria for TMS were applied. The study was approved by the local ethics committee of [Authors Region] (protocol number 2009-A01087-50) and followed the Code of Ethics of the World Medical Association (Declaration of Helsinki).

#### Stimulus procedure

Participants were placed 57 cm from the screen (36.5° x 27° of visual angle) in a dark room. Their head was maintained by a chinrest and headrest. They performed 26 blocks of 72 trials each. One block was used for practice. One block allowed the determination of the Stimulus Onset Asynchrony (SOA) to reach approximately 70% correct, using a staircase procedure. Then, 24 blocks corresponded to the main experiment: 4 blocks with no TMS, 10 blocks with TMS applied over the FEF, and 10 blocks applied over the vertex (control; see *TMS procedure*).

Participants performed a difficult visual search (**Figure 1**; same procedure as in Dugué et al., 2011; 2015a): report the presence or absence of a target letter T, among distractor letters Ls (1.5° x 1.5°). On each trial, four stimuli were presented on the left hemifield at constant eccentricity (6°): either four Ls (target absent trials) or three Ls and one T (target present trials), randomly presented in four orientations (0, 90, 180 or 270° from upright). Stimuli were always presented in the left visual field. Accuracy, as per dprime, was our main dependent variable. Thus, participants were asked to respond accurately, and with no time pressure, by pressing a key on the keyboard.

**Figure 1:**
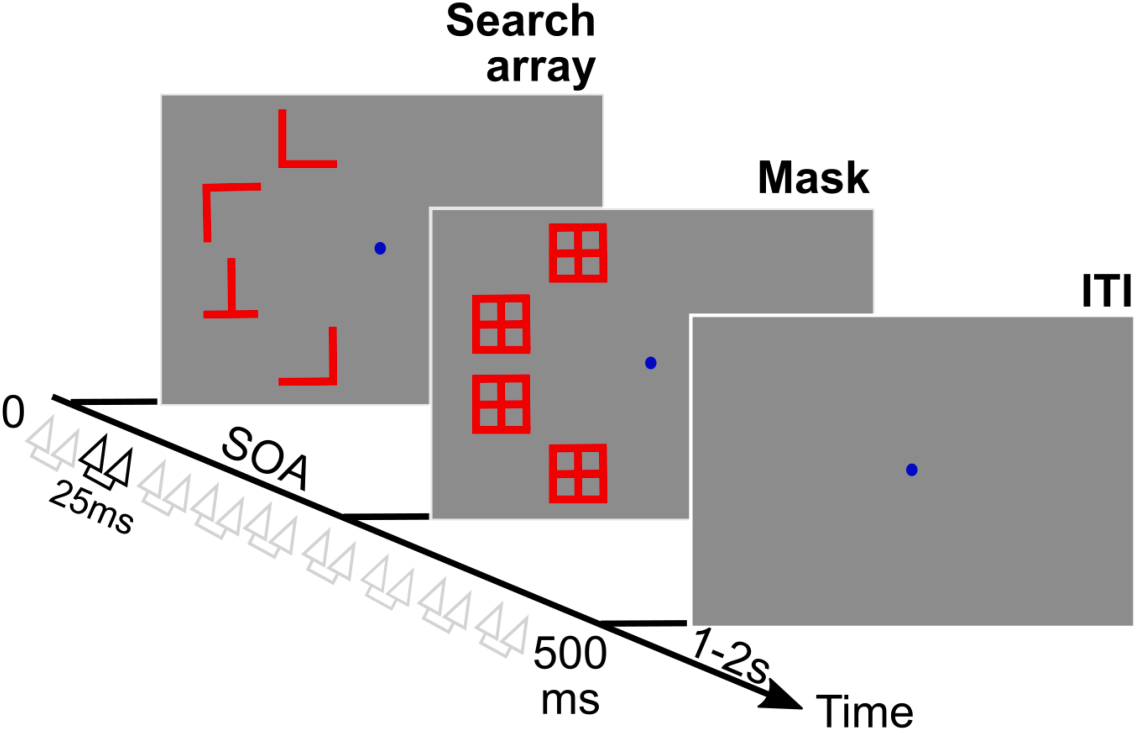
Experimental protocol. While participants performed a visual search (finding a T letter among Ls), they were stimulated over the right-FEF or the vertex (control) with a double-pulse of TMS (25 ms interval) applied at random delays between 50 ms and 450 ms (50 ms increments) after the search array onset.

The Stimulus Onset Asynchrony (SOA), i.e. the delay between search array onset and mask onset, was predefined for each observer to achieve about 70% correct (85 ms ± 8 ms). The total trial duration (including search array and masks) was 500 ms for all participants.

#### TMS targeting and delivery

TMS pulses were delivered using a 70-mm figure-of-eight coil (biphasic stimulator, Magstim Rapid^2^). A structural T1-weighted MRI scan (3T Philips, flip angle = 8°, TR = 8.1 ms, TE = 3.7 ms, FOV = 240 x 240 mm, voxel size = 1mm isotropic) was acquired for 11 participants at the imagery platform of the [Author University]. For these participants, the right FEF was localized on each individual MRI using averaged Talairach coordinates x = 31, y = −2, z = 47 (Paus, 1996), and a 0.5 radius spherical region of interest (ROI; same procedure as in Chanes et al., 2012; 2013). The final MRI was uploaded into a frameless stereotaxic system and reconstructed in 3D for its use in an online TMS neuronavigation system (eXimia NBS system, Nextim).

Participants were all wearing an EEG cap to help localize the stimulated ROI on the surface. Eight participants did not have anatomical MRI. For those participants, the stimulation ROI was determined as the barycenter of the region of stimulation from the 11 previous participants.

The TMS coil was placed tangentially to the skull and its handle oriented 45° in a rostral-to-caudal and lateral-to-medial orientation. The stimulation intensity started at 50% of the TMS machine maximal output, and was then adjusted just below the threshold of facial and temporal muscle activation (average intensity across participants = 52% ± 2% SEM). For comparison, (Chanes et al., 2012) applied single pulses of TMS at ∼67% of the TMS machine maximum output, whereas (Chanes et al., 2013) applied a train of four pulses of TMS at 30 Hz at ∼44%.

The Vertex was used as a stimulation control site for non-specific TMS effect such as clicking noise and tapping sensation. This region was localized for each participant as the region under electrode Cz on the EEG cap (O’Shea et al., 2004).

#### TMS procedure

Double-pulses of TMS (25 ms interval) were applied at random delays after search array onset (9 possible delays, from 50 to 450 ms, 50 ms increments; see **Figure 1**). Right-FEF and Vertex stimulations were blocked. Half of the participants (randomly assigned) performed the right-FEF blocks first, while the other half started with the vertex ones. Participants performed 80 trials per stimulation delay and condition.

#### Re-analysis of two previously published datasets

In the current studies, we compare the effect of TMS applied at various delays over the FEF during a difficult visual search task, with the results of two previously published studies using the same search task (L vs. T), while TMS was applied over the occipital pole (V1/V2).

The first dataset comes from the published study by (Dugué et al., 2011). Based on phosphene mapping, double-pulses of TMS (25 ms interval) were applied at one of various delays (8 possible delays from 100 to 450 ms, 50 ms increments) to a consistent brain location in retinotopic areas (V1/V2). The search array was presented either at the location affected by the TMS pulses (phosphene region) or in the symmetric region in the opposite hemifield (retinotopically-defined control region). Thus, the stimulation was identical over the cortex but was either interfering with the stimulus, retinotopic location (phosphene condition), or not (control condition; see Dugué et al., 2011, for further methodological details).

The second dataset comes from the published study by (Dugué et al., 2015a). In this study, the authors followed the same procedure as in (Dugué et al., 2011). The only difference is the way the two pulses were administered. In each trial, one pulse remained fixed at a latency of 312.5 ms after the search array onset, based on the main effect found previously in (Dugué et al., 2011). The second pulse was applied at 13 other possible delays before or after the first pulse (112.5, 137.5, 162.5, 187.5, 212.5, 237.5, 262.5, 287.5, 337.5, 362.5, 387.5, 412.5, or 437.5 ms after stimulus onset; see Dugué et al., 2015a, for further methodological details).

#### Fourier Analysis

For all three studies, we calculated a dprime modulation index as the main dependent variable by subtracting the main stimulation condition and the control condition. In the current study, dprime modulation was the difference between the right-FEF and the vertex condition trials. In the two previously published V1 studies, dprime modulation was the difference between the phosphene condition and the control condition trials (see previous section). In other words, in all three cases, negative values corresponded to a target region-specific impairment of performance by TMS.

We first combined the results from all three studies to investigate the overall TMS modulation of attentional performance during the difficult search task. Since the TMS pulses were not applied at the same delays across the three studies, we first oversampled each individual dprime modulation time-course every 12.5 ms using a linear interpolation. We then averaged all three datasets together (see **Figure 2A**). On this pooled dataset, we performed a Fast Fourier Transform (FFT performed on actual time points measured, i.e. no oversampling; see **Figure 2B**). Bootstrapping assessed the significance of each frequency component: The simulations were obtained by shuffling the labels of TMS delays, following the null hypothesis that the dprime modulation was independent of TMS latency (100,000 iterations). The 100,000 surrogate amplitude spectra were ranked in ascending order, separately for each delay. The 95,001th value was considered as the limit of the 95% ci (p < 0.05). To take into consideration multiple comparisons, an experimentally observed spectral amplitude value was considered significantly different from the corresponding null distribution when p < 0.001 (99.9% ci).

**Figure 2:**
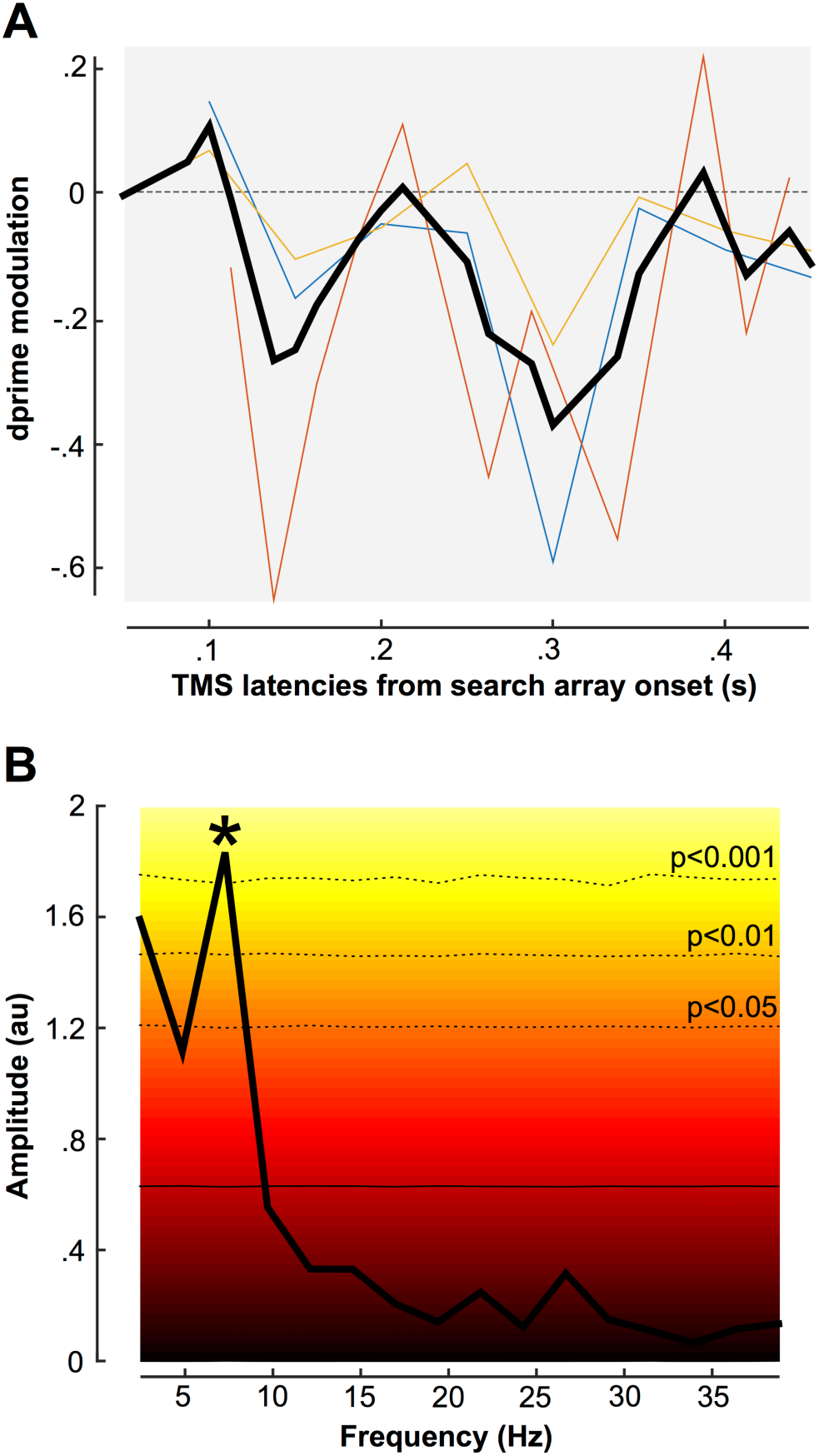
TMS modulates attentional search periodically. (A) Dprime modulations (test – control condition) are represented as a function of TMS latencies from the search array onset. The color lines represent each individual study (yellow: current FEF study; blue: first V1 study (Dugué et al., 2011); red: second V1 study (Dugué et al., 2015a). The black line is the average across all three studies. **(B)** Amplitude spectrum obtained by FFT decomposition of the averaged data across the three studies. The background color represents the level of significance obtained with a bootstrap procedure. The bottom, horizontal black line represents the amplitude spectrum of the surrogate distribution. The dashed lines represent different levels of significance. * symbol represents the significant spectral component at 7 Hz (p<0.001).

We then looked at the amplitude spectra of each single study using FFT decomposition. On these individual datasets, we performed an FFT on data padded (using the average value) to get a 4000 ms segment. Note that we also did the analysis on non-padded data and obtained comparable results. Here, the significance of each oscillatory component was assessed by nonparametric statistics. Monte Carlo simulations were performed under the null hypothesis that the dprime modulation was independent of TMS latency (100,000 iterations). For each iteration, we recomputed the grand-averaged curve of the difference of dprime between test and control conditions, and its amplitude spectrum. For each oscillatory frequency, we then sorted these surrogates in ascending order and calculated confidence intervals and the corresponding p-values (see **Figure 3**, left column). A frequency peak was considered significantly different from the corresponding null distribution with p < 0.001.

**Figure 3:**
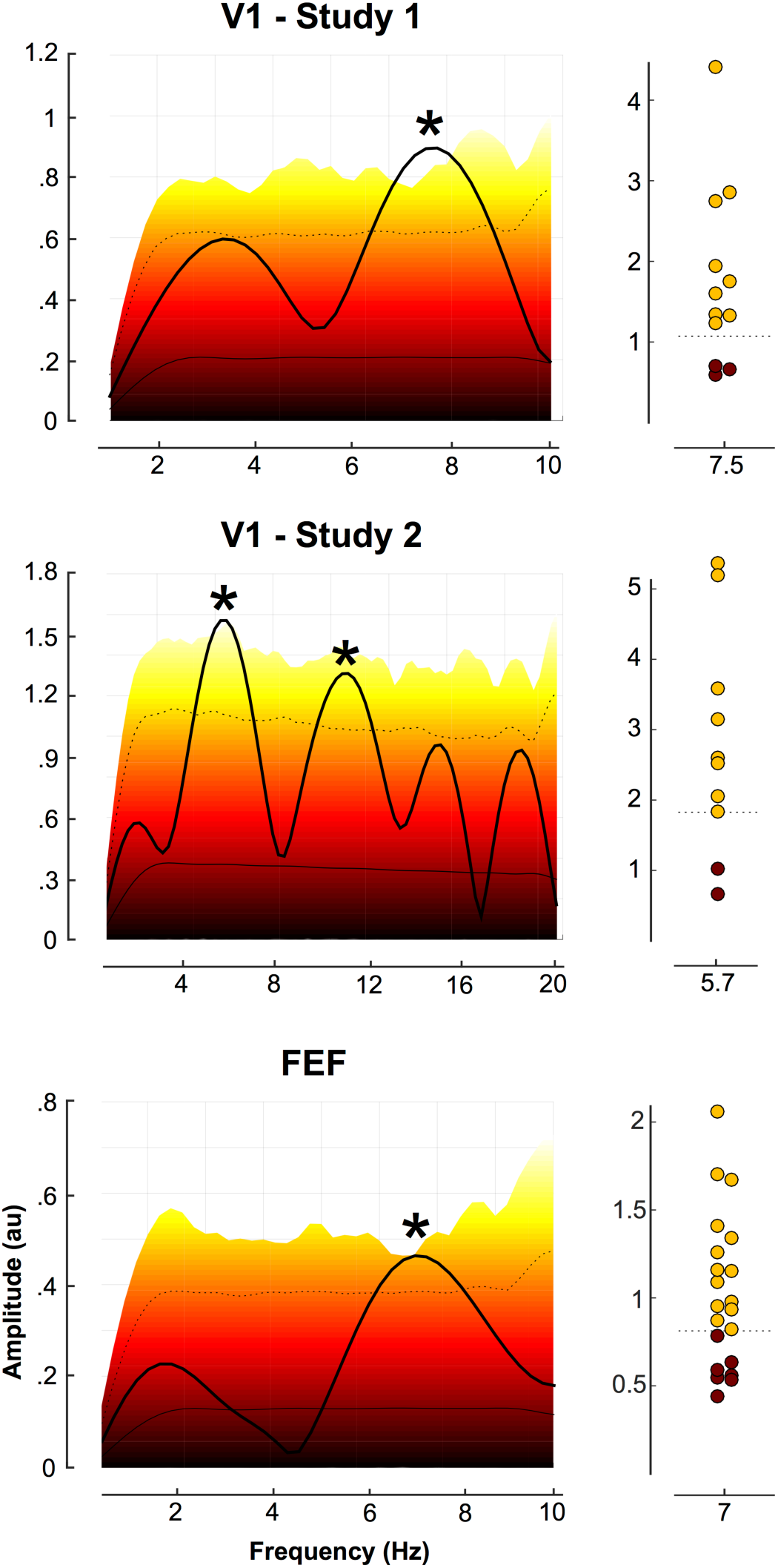
Attentional periodicity in V1 and FEF. For each study, the graphs in the left column represent the amplitude spectra obtained by FFT decomposition of the averaged performance (as per dprime modulation; see Methods) across participants. Note the distinct frequency axis in the middle panel, due to the increased time resolution (and corresponding Nyquist frequency) in that study. The background color represents the level of significance obtained with a Monte Carlo procedure. The bottom, horizontal black line represents the amplitude spectrum of the surrogate distribution. The dashed line represents p < 0.001. * symbols represent the significant spectral components. The right column represents the amplitude of the significant spectral components (of the left column) for each participant. The dashed line represents p<0.001 (the yellow dots correspond to participants for which the amplitude passes this level of significance).

Finally, for each participant in each study, we looked at the significant frequency peak observed in the amplitude spectra obtained by frequency decomposition of the averaged dprime modulation across participants. Similarly as before, using Monte Carlo simulations we evaluated whether this frequency component was significantly different from the corresponding null distribution (with p < 0.001) for each participant (see **Figure 3**, right column).

## Results

To test the prediction that difficult visual search periodically involves low- and high-level regions along with iterative attentional selection, we conducted a TMS experiment in which we interfere with the FEF at various delays while observers performed a task in which they have to report the presence or absence of the letter T among letter Ls. Dprime modulation was calculated as the difference between the right-FEF and the vertex (control) stimulation. These results were first combined with the results of two previously published studies (Dugué et al., 2011; 2015a) using the same search task (L vs. T), while TMS was applied either over the retinotopic location of the early visual cortex corresponding to the search array location, or over the symmetric (control) location (see Materials and Methods). Note that a meta-analysis of the literature on TMS studies of attention during visual search reveals that no other study had the necessary temporal sampling resolution for such an investigation (i.e. single or double-pulses of TMS sampling a large time window at multiple delays on separate trials; see **Table 1**). **Figure 2A** represents the combined dprime modulation across all three studies, as a function of the delays at which TMS was applied during the difficult search task.

We further investigated the temporal dynamics of such performance modulation. An FFT applied to this combined dataset revealed a significant peak at 7 Hz suggesting that TMS periodically interferes with search performance at the theta frequency (**Figure 2B**).

Critically, to understand the origin of this overall, descriptive effect, we performed the same frequency decomposition on each individual dataset (**Figure 3**). In all three studies, we observed a significant peak in the theta frequency range: 7.5 Hz for the first V1 study (Dugué et al., 2011), 5.7 Hz for the second V1 study (Dugué et al., 2015a), and 7 Hz for the current, FEF study. Importantly, this peak was observed in most of the observers (**Figure 3,** right column) suggesting that the periodicity is not a mere effect of performance averaging.

## Discussion

Using TMS applied at various delays while observers performed a difficult, attentional search task, we showed that both V1 and the FEF are involved periodically during the search, at the theta frequency (∼6-7 Hz). This finding supports the idea that visual search tasks are processed by a hierarchical system involving periodic, iterative connections between low- and high-level regions until target recognition. This hypothesis is in line with the large accumulation of evidence that attention acts via feedback to sensory areas (Motter, 1994; Mehta et al., 2000; Schroeder et al., 2001; Kastner and Pinsk, 2004; Bressler et al., 2008).

An additional prediction made by such hierarchical two-way processing stream is that if both the low- and high-level regions are periodically sending information to each other, there should also be a phase lag between their respective modulations. Unfortunately, because the results were obtained from independent studies (different participants and sample sizes) and the peak frequency was not the same across the three studies, we were not able to compare their phases. In the future, one could perform an experiment in which the same observers are stimulated at various delays over the FEF and V1, while performing the same difficult search task. This would allow the characterization within the same participants of the respective temporal dynamics of V1 and FEF, and their interaction, during difficult visual search.

Oscillations in behavioral performance have been the topic of a large, recent body of research. Two rhythm frequencies have been reported (VanRullen, 2016), i.e. alpha (∼10 Hz) and theta (∼7Hz). It has been suggested that while the alpha rhythm reflects an intrinsic, sensory rhythm, sampling information at a single location, theta rather reflects attentional exploration, sampling information at multiple locations (Dugué and VanRullen, 2017). This hypothesis is in line with a recent experiment in which attentional exploration was explicitly manipulated using a cueing paradigm (Dugué et al., 2016). By applying TMS at various delays over V1, the authors demonstrated that performance in a 2-AFC orientation discrimination task was modulated by TMS periodically at the theta frequency (∼5 Hz) only when attention had to be reallocated from a distractor to a target location.

One might wonder whether the observed periodicity in all the previously described TMS studies (including the present one) is due to a true, intrinsic property of the attention system, which processes information periodically, or whether it is actually induced by the TMS. One critical piece of evidence in favor of the former is that in (Dugué et al., 2015a) the authors not only observed a periodicity in behavioral performance due to the stimulation, but also showed in independent trials without TMS (but in the same participants) that brain oscillations (as measured by EEG) at the same frequency (∼6 Hz; theta) correlated with search performance. Consequently, oscillations likely reflect a periodicity in cortical excitability, and TMS is thus able to probe the system at different excitability states.

In the present study, we show that the FEF is involved at the theta frequency during this attentional search. Previous studies investigating the spontaneous activity of the fronto-parietal region (Rosanova et al., 2009), and the role of the FEF in attentional search in monkeys (Buschman and Miller, 2009) and humans (Phillips and Takeda, 2010) however showed periodicity in the low (13-24 Hz) and high beta frequency range (36-56 Hz). Given the use of multiple delays in the different TMS studies presented here, frequencies above 10 Hz could not be characterized. Thus, we cannot rule out that other, higher frequencies are related to attentional sampling during this difficult visual search.

In the present study, we investigated the temporal dynamics of V1 and FEF during attentional search, and revealed that both regions are involved periodically at the theta frequency. This study brings convincing, converging evidence, together with multiple studies using various approach including psychophysics, EEG and TMS, and analysis tools, in favor of a theta, intrinsic rhythm as the support of attentional exploration.

## Abbreviated Title

FEF and attentional periodicity

## Author Contributions

LD and RV conceived and supervised the study. LD and RV designed the experiment. LD, AAB and PM carried out the experiment. LD, AAB and RV analyzed the data. LD wrote the first draft of the manuscript. LD, AAB, PM and RV finalized the manuscript and approved the final version of the manuscript.

## Acknowledgements

The authors thank Samuel Planton for his help with the FEF localization based on structural MRI.

## References

Ashbridge E (1997) Temporal aspects of visual search studied by transcranial magnetic stimulation. Neuropsychologia 35:1121–1131.

Bressler SL, Tang W, Sylvester CM, Shulman GL, Corbetta M (2008) Top-Down Control of Human Visual Cortex by Frontal and Parietal Cortex in Anticipatory Visual Spatial Attention. J Neurosci 28:10056–10061.

Busch NA, VanRullen R (2010) Spontaneous EEG oscillations reveal periodic sampling of visual attention. Proc Natl Acad Sci USA 107:16048–16053.

Buschman TJ, Miller EK (2009) Serial, Covert Shifts of Attention during Visual Search Are Reflected by the Frontal Eye Fields and Correlated with Population Oscillations. Neuron 63:386–396.

Chanes L, Chica AB, Quentin R, Valero-Cabré A (2012) Manipulation of pre-target activity on the right frontal eye field enhances conscious visual perception in humans. Geng JJ, ed. PLoS ONE 7:e36232.

Chanes L, Quentin R, Tallon-Baudry C, Valero-Cabré A (2013) Causal frequency-specific contributions of frontal spatiotemporal patterns induced by non-invasive neurostimulation to human visual performance. Journal of Neuroscience 33:5000–5005.

Corbetta M, Shulman GL (2002) Control of goal-directed and stimulus-driven attention in the brain. Nat Rev Neurosci 3:201–215.

Deco G, Pollatos O, Zihl J (2002) The time course of selective visual attention: theory and experiments. Vision Res 42:2925–2945.

Dugué L, Marque P, VanRullen R (2011) Transcranial magnetic stimulation reveals attentional feedback to area V1 during serial visual search. Herzog MH, ed. PLoS ONE 6:e19712.

Dugué L, Marque P, VanRullen R (2015a) Theta oscillations modulate attentional search performance periodically. J Cogn Neurosci 27:945–958.

Dugué L, McLelland D, Lajous M, VanRullen R (2015b) Attention searches nonuniformly in space and in time. Proc Natl Acad Sci USA 112:15214–15219.

Dugué L, Roberts M, Carrasco M (2016) Attention Reorients Periodically. Current Biology 26:1595–1601.

Dugué L, VanRullen R (2014) The dynamics of attentional sampling during visual search revealed by Fourier analysis of periodic noise interference. Journal of Vision 14:11–11.

Dugué L, VanRullen R (2017) Transcranial Magnetic Stimulation Reveals Intrinsic Perceptual and Attentional Rhythms. Front Neurosci 11:154.

Dugué L, Xue AM, Carrasco M (2017) Distinct perceptual rhythms for feature and conjunction searches. Journal of Vision 17:22–22.

Eckstein MP (2011) Visual search: A retrospective. Journal of Vision 11:14–14.

Ellison A, Lane AR, Schenk T (2007) The interaction of brain regions during visual search processing as revealed by transcranial magnetic stimulation. Cereb Cortex 17:2579–2584.

Ellison A, Schindler I, Pattison LL, Milner AD (2004) An exploration of the role of the superior temporal gyrus in visual search and spatial perception using TMS. Brain 127:2307–2315.

Fiebelkorn IC, Pinsk MA, Kastner S (2018) A Dynamic Interplay within the Frontoparietal Network Underlies Rhythmic Spatial Attention. Neuron 99:842–853.e848.

Fiebelkorn IC, Saalmann YB, Kastner S (2013) Rhythmic Sampling within and between Objects despite Sustained Attention at a Cued Location. Current Biology 23:2553–2558.

Fuggetta G, Pavone EF, Walsh V, Kiss M, Eimer M (2006) Cortico-cortical interactions in spatial attention: A combined ERP/TMS study. J Neurophysiol 95:3277–3280.

Helfrich RF, Fiebelkorn IC, Szczepanski SM, Lin JJ, Parvizi J, Knight RT, Kastner S (2018) Neural Mechanisms of Sustained Attention Are Rhythmic. Neuron 99:854–865.e855.

Huang Y, Chen L, Luo H (2015) Behavioral oscillation in priming: competing perceptual predictions conveyed in alternating theta-band rhythms. Journal of Neuroscience 35:2830–2837.

Itti L, Koch C (2001) Computational modelling of visual attention. Nat Rev Neurosci 2:194–203.

Juan C-H, Walsh V (2003) Feedback to V1: a reverse hierarchy in vision. Exp Brain Res 150:259– 263.

Kalla R, Muggleton NG, Cowey A, Walsh V (2009) Human dorsolateral prefrontal cortex is involved in visual search for conjunctions but not features: A theta TMS study. Cortex 45:1085–1090.

Kalla R, Muggleton NG, Juan C-H, Cowey A, Walsh V (2008) The timing of the involvement of the frontal eye fields and posterior parietal cortex in visual search. NeuroReport 19:1067–1071.

Kastner S, Pinsk MA (2004) Visual attention as a multilevel selection process. Cognitive, Affective, & Behavioral Neuroscience 4:483–500.

Kastner S, Pinsk MA, De Weerd P, Desimone R, Ungerleider LG (1999) Increased Activity in Human Visual Cortex during Directed Attention in the Absence of Visual Stimulation. Neuron 22:751–761.

Landau AN, Fries P (2012) Attention Samples Stimuli Rhythmically. Current Biology 22:1000–1004.

Landau AN, Schreyer HM, van Pelt S, Fries P (2015) Distributed Attention Is Implemented through Theta-Rhythmic Gamma Modulation. Current Biology 25:2332–2337.

Lane AR, Smith DT, Schenk T, Ellison A (2011) The involvement of posterior parietal cortex in feature and conjunction visuomotor search. J Cogn Neurosci 23:1964–1972.

Mehta AD, Ulbert I, Schroeder CE (2000) Intermodal Selective Attention in Monkeys. II: Physiological Mechanisms of Modulation. Cereb Cortex 10:359–370.

Motter BC (1994) Neural correlates of attentive selection for color or luminance in extrastriate area V4. J Neurosci 14:2178–2189.

Muggleton NG, Cowey A, Walsh V (2008) The role of the angular gyrus in visual conjunction search investigated using signal detection analysis and transcranial magnetic stimulation. Neuropsychologia 46:2198–2202.

Muggleton NG, Juan C-H, Cowey A, Walsh V (2003) Human Frontal Eye Fields and Visual Search. J Neurophysiol 89:3340–3343.

Muggleton NG, Kalla R, Juan C-H, Walsh V (2011) Dissociating the contributions of human frontal eye fields and posterior parietal cortex to visual search. J Neurophysiol 105:2891–2896.

Nakayama K, Martini P (2011) Situating visual search. Vision Res 51:1526–1537.

O’Shea J, Muggleton NG, Cowey A, Walsh V (2004) Timing of Target Discrimination in Human Frontal Eye Fields. Journal of Cognitive Neuroscience 16:1060–1067.

Palmer J, Ames CT, Lindsey DT (1993) Measuring the effect of attention on simple visual search. Journal of Experimental Psychology: Human Perception and Performance 19:108–130.

Paus T (1996) Location and function of the human frontal eye-field: a selective review. Neuropsychologia 34:475–483.

Phillips S, Takeda Y (2010) Frontal-parietal synchrony in elderly EEG for visual search. Int J Psychophysiol 75:39–43.

Rosanova M, Casali A, Bellina V, Resta F, Mariotti M, Massimini M (2009) Natural Frequencies of Human Corticothalamic Circuits. J Neurosci 29:7679–7685.

Saalmann YB, Pigarev IN, Vidyasagar TR (2007) Neural mechanisms of visual attention: how top-down feedback highlights relevant locations. Science 316:1612–1615.

Schindler I, Ellison A, Milner AD (2008) Contralateral visual search deficits following TMS. J Neuropsychol 2:501–508.

Schroeder CE, Mehta AD, Foxe JJ (2001) Determinants and mechanisms of attentional modulation of neural processing. Front Biosci.

Song K, Meng M, Chen L, Zhou K, Luo H (2014) Behavioral oscillations in attention: rhythmic α pulses mediated through θ band. Journal of Neuroscience 34:4837–4844.

Treisman A (1998) Feature binding, attention and object perception. Philosophical Transactions of the Royal Society of London B: Biological Sciences 353:1295–1306.

van Diepen RM, Miller LM,Mazaheri A, Geng JJ(2016) The Role of Alpha Activity in Spatial and Feature-Based Attention. eNeuro 3:ENEURO.0204–16.2016.

VanRullen R (2013) Visual Attention: A Rhythmic Process? Current Biology 23:R1110–R1112.

VanRullen R (2016) Perceptual Cycles. Trends Cogn Sci (Regul Ed) 20:723–735.

VanRullen R, Carlson T, Cavanagh P (2007) The blinking spotlight of attention. Proc Natl Acad Sci USA 104:19204–19209.

Walsh V, Ashbridge E, Cowey A (1998) Cortical plasticity in perceptual learning demonstrated by transcranial magnetic stimulation. Neuropsychologia 36:363–367.

Yan Y, Wei R, Zhang Q, Jin Z, Li L (2016) Differential roles of the dorsal prefrontal and posterior parietal cortices in visual search: a TMS study. Sci Rep 6:30300.

